# High efficacy of therapeutic equine hyperimmune antibodies against SARS-CoV-2 variants of concern

**DOI:** 10.1101/2021.06.12.448080

**Authors:** Andres Moreira-Soto, Mauricio Arguedas, Hebleen Brenes, Willem Buján, Eugenia Corrales-Aguilar, Cecilia Díaz, Ann Echeverri, Marietta Flores-Díaz, Aarón Gómez, Andrés Hernández, María Herrera, Guillermo León, Román Macaya, Arne Kühne, José Arturo Molina-Mora, Javier Mora, Alfredo Sanabria, Andrés Sánchez, Laura Sánchez, Álvaro Segura, Eduardo Segura, Daniela Solano, Claudio Soto, Jennifer L. Stynoski, Mariángela Vargas, Mauren Villalta, Chantal B E M Reusken, Christian Drosten, José María Gutiérrez, Alberto Alape-Girón, Jan Felix Drexler

**Author notes:** Corresponding author **Corresponding author contact information** Professor Dr. Jan Felix Drexler, Helmut-Ruska-Haus, Institute of Virology, Campus Charité Mitte, Charitéplatz 1, 10098 Berlin, Germany, /Fax. **Competing Financial Interests Statement** The authors declare no competing financial interests.

## Abstract

SARS-CoV-2 variants of concern (VoC) show reduced neutralization by vaccine-induced and therapeutic monoclonal antibodies. We tested therapeutic equine polyclonal antibodies (pAbs) against four VoC (alpha, beta, epsilon and gamma). We show that equine pAbs efficiently neutralize VoC, suggesting they are an effective, broad coverage, low-cost and a scalable COVID-19 treatment.

## To the editor

SARS-CoV-2 causes coronavirus infectious disease 19 (COVID-19), which leads to either critical illness or death in 5% of patients (1). COVID-19 prevention and treatment options include vaccines, antivirals, and antibody formulations. A wide array of vaccine platforms have shown efficacy in preventing severe disease, but universal access is limited and many resource-limited settings largely lack sufficient vaccine coverage (2). Even though there are more than 300 therapeutic drugs in clinical trials, few have proven advantageous, such as dexamethasone (1, 3). Direct-acting antivirals like Remdesivir are most effective if given very early, require supplementary oxygen therapy and are very costly at 2,000-3,000 USD per treatment, limiting universal access (4). Convalescent plasma or hyperimmune globulins, which can be prepared from the pooling of many donors, have been used for decades to treat diseases such as ebola and influenza and could be a more affordable at 350-1,000 USD per treatment. However, their preparation is donor-dependent, requires strict donor rigorous testing for both blood-borne pathogens and high levels of neutralizing anti-SARS-CoV-2 antibodies, not readily available on blood bank systems in many developing countries (5). The use of monoclonal antibodies (mAbs) are safe alternatives shown to enhance viral clearance, but their large scale production is challenging and costly, at around 1,500-6,500 USD per treatment (6). A low-cost alternative to mAbs are formulations of intact or fragmented equine polyclonal antibodies (pAbs), widely used for decades as therapies against viral infections or as antivenoms.

We and others have previously shown that horses can be efficiently immunized with different SARS-CoV-2 antigens to yield high quantities of purified polyclonal antibodies (pAbs) that are 50-80 times more potent than convalescent plasma (7, 8). A formulation of equine polyclonal F(ab’)_2_ fragments against the receptor binding domain (RBD) of SARS-CoV-2 was tested in a multi-center, double-blind, placebo-controlled phase II/III clinical trial showing that it is well tolerated and leads to clinical improvement of hospitalized patients with moderate to severe COVID-19 (9). Additionally, there is an ongoing randomized, multi-center, double-blind, placebo-controlled, dose-finding, phase IIb/III clinical trial (NCT04838821) at hospitals of the Costa Rican Social Security Fund testing equine pAbs formulations to treat moderate and severe COVID-19 cases.

However, pre-clinical data of equine hyperimmune pAbs are only available for early SARS-CoV-2 isolates, such data are lacking for recent and globally circulating variants, considered of concern (VoC) due to their increased transmissibility. Voc alpha, beta, epsilon and gamma (https://www.cdc.gov/coronavirus/2019-ncov/variants/variant-info.html) (lineage designations in Pango/Nextrain: B.1.1.7/501Y.V1 first detected in the United Kingdom, B.1.351/501Y.V2 first detected in South Africa, P.1/501Y.V3 first detected in Brazil/Japan, and B.1.427/B.1.429 first detected in the US/California) exhibit a substantial reduction or complete abrogation of neutralization by therapeutic mAbs or by antibodies present in the plasma of vaccinated or convalescent individuals (10).

Here we report the results of a plaque reduction neutralization assay against VoC for our purified equine pAbs formulations. The two formulations are the SARS-CoV-2 recombinant S1 protein (called anti-S1; produced in baculovirus insect cells), and SEM mosaic (called anti-mix; an *E. coli* derived recombinant protein containing the S, E, and M immunodominant regions) derived from the strain Wuhan-Hu-1, Accession N YP_009724390 (Native Antigen Company, Oxford, United Kingdom), purified using caprylic acid precipitation method (8). Both formulations effectively neutralized four VoC and an early isolate of the virus (Germany/Gisaid_EPI_ISL_406862) at similar inhibitory concentrations (IC_50_ range for anti-S1 formulation: 0.206-0.377 μg/mL; and for the anti-mix formulation: 0.146-0.471 μg/mL; **Figure 1**; IC_50_ dose-response curves are shown in the Technical Annex). Those concentrations are extremely low when compared to pAbs doses used by other groups in patients in clinical trials (4 mg/kg) (9), even at the upper estimates of the 95% confidence intervals, reaching a maximum of 13.89 μg/mL for the beta (B.1.351/501Y.V2) VoC. For both equine pAbs formulations the differences between potencies against tested VoC and early SARS-CoV-2 isolates were not statistically significant (sum-of-squares F test of Anti-S1; p=0.9, Anti-Mix, p=0.8).

**Figure 1.**
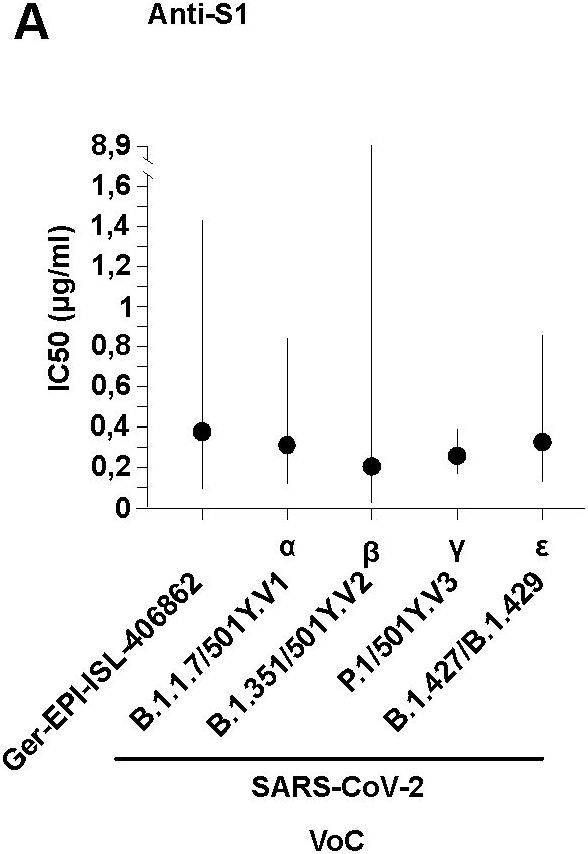
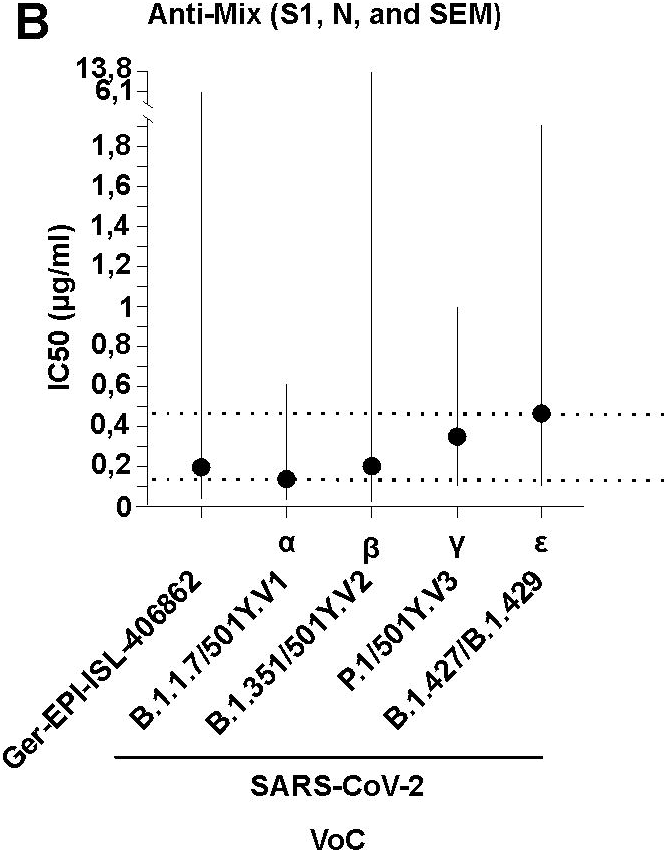
*In vitro* neutralizing potency of (A) Anti-S1 (S1 SARS-CoV-2 recombinant protein) and (B) Anti-Mix (mixture of S1, N, and SEM mosaic SARS-CoV-2 recombinant proteins of Wuhan-Hu-1, Accession N YP_009724390.1) polyclonal antibodies purified from the plasma of hyperimmunized horses against different SARS-CoV-2 variants of concern (VoC) and a early isolate, named using WHO and Pango/Nextrain designations (strains used= GERMANY/GISAID EPI_ISL 406862, BetaCoV/ChVir21652, hCoV-19/Aruba_11401/2021, hCoV-19/Netherlands/NoordHolland_10915/2021, BetaCoV/ChVir22131/B.1.351/501Y.V2, acquired from https://www.european-virus-archive.com/evag-news/sars-cov-2-collection). The inhibitory concentration (IC_50_) in plaque reduction neutralization tests (PRNT) was calculated using a nonlinear regression analysis in the GraphPadPrism 5 software. Potencies (IC_50_) were not statistically different among viral variants with either formulation, and the null hypothesis was not rejected, meaning the IC50 was equal in all datasets. Dotted lines denote the mean minimum and maximum concentration and solid lines denote 95% confidence intervals for both formulations. Plaque reduction neutralization tests (PRNT) were performed as follows. Briefly, VeroE6 cells (3.25 × 105 cells/ml) were seeded in 24-well plates and incubated overnight. Equine polyclonal antibody formulations were mixed in equal parts with a virus solution containing 20 PFU. The serum–virus solution was incubated at 37°C for 1 h and added to the cells. After 1 h at 37°C, supernatants were discarded, and cells were supplemented with 1.2% Avicel solution in DMEM. After 3 d at 37°C, supernatants were removed, and the 24-well plates were fixed and inactivated using a 6% formaldehyde/PBS solution and stained with crystal violet, and plaques were counted.

Our data suggest high potential of equine pAbs for treatment of COVID-19. By shifting antivenom platforms to produce equine pAbs, laboratories in both developed and developing countries that have been manufacturing and distributing safe and standardized antivenoms for decades could rapidly fill the gaps in global demand for therapies that are both effective against VoC and affordable to low- and middle-income countries.

## Supporting information

Anex Figure

## Acknowledgements

This work was supported by the “Global Centres for Health and Pandemic Prevention” from the German academic exchange services (DAAD).

**Annex figure:** IC50 dose-response curves to SARS-CoV-2 early isolates and variants of concern named using WHO and Pango/Nextrain designations. The Y axis denotes the mean plaque forming units (PFU) per milliliter in triplicate. The X axis denotes the Log10 concentration of the Anti-S1 and the Anti-Mix (combination of S1, N and SEM mosaic protein of Wuhan-Hu-1, Accession N YP_009724390.1) formulations.

## First author biographical sketch

Dr. Moreira-Soto is a virologist at the Charité-Universitätsmedizin Berlin Institute of Virology and the University of Costa Rica Research Center for Tropical Diseases (CIET). His research interests include virology, epidemiology, public health, and molecular biology of emerging infectious diseases.

