## Supplementary figures and images for "High efficacy of therapeutic equine hyperimmune antibodies against SARS-CoV-2 variants of concern"

### Anex Figure

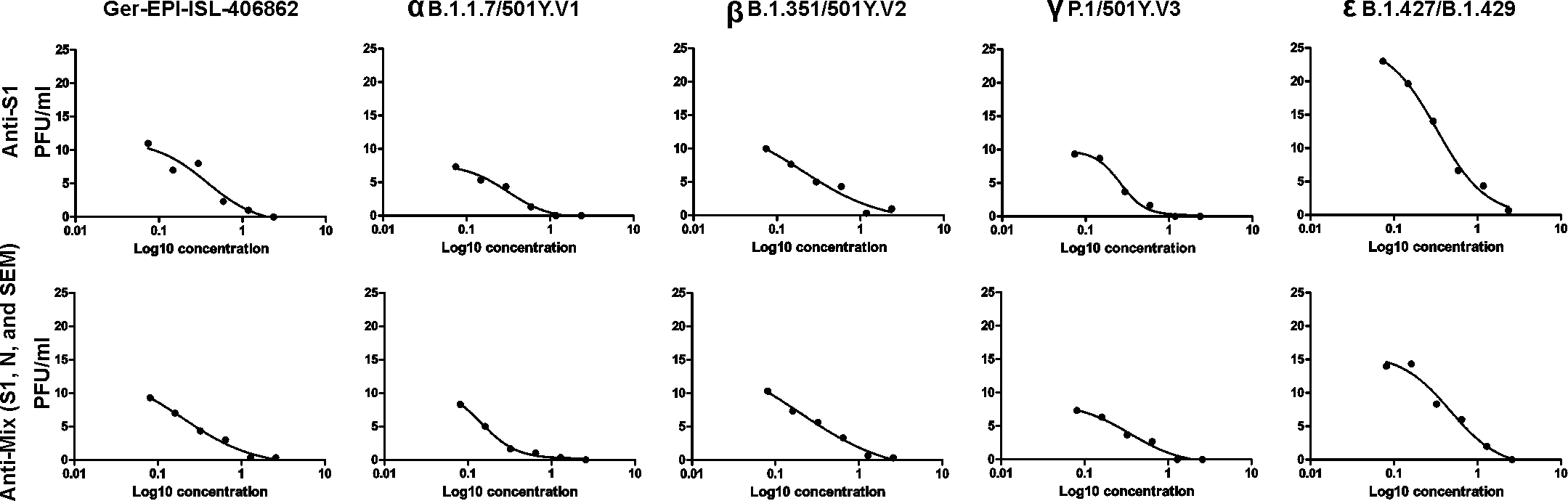
